# Supervised deep machine learning models predict forelimb movement from excitatory neuronal ensembles and suggest distinct pattern of activity in CFA and RFA networks

**DOI:** 10.1101/2024.01.30.577967

**Authors:** Shahrzad Latifi, Jonathan Chang, Mehdi Pedram, Roshanak Latifikhereshki, S Thomas Carmichael

**Author notes:** Corresponding authors: Shahrzad Latifi, S Thomas Carmichael.

## Abstract

Neuronal networks in the motor cortex are crucial for driving complex movements. Yet it remains unclear whether distinct neuronal populations in motor cortical subregions encode complex movements. Using *in vivo* two-photon calcium imaging (2P) on head-fixed grid-walking animals, we tracked the activity of excitatory neuronal networks in layer 2/3 of caudal forelimb area (CFA) and rostral forelimb area (RFA) in motor cortex. Employing supervised deep machine learning models, a support vector machine (SVM) and feed forward deep neural networks (FFDNN), we were able to decode the complex grid-walking movement at the level of excitatory neuronal ensembles. This study indicates significant differences between RFA and CFA decoding accuracy in both models. Our data demonstrate distinct temporal-delay decoding patterns for movements in CFA and RFA, as well as a selective ensemble of movement responsive neurons with higher distribution in CFA, suggesting specific patterns of activity-induced movement in these two networks.

## Introduction

The brain is network of highly optimized, connected systems across multiple spatiotemporal scales. The information pathways within these specialized multi-scale networks, from synapses to brain regions, enable motion, cognition, and perception. Elucidating how these functions derive from brain activity is one of the fundamental goals in neuroscience and thus decoding neuronal activities that read out complex behaviors such as cognitive processes or motor function has become a pivotal analytic tool, but still challenging. Initially, traditional decoding methods such as multiple linear regressions or nonlinear dynamic models have been used to predict behaviors such as animal’s movement trajectories based on extracted signals from motor cortex or spatial locations associated with hippocampal neuronal activity [1, 2]. Because of the complexity of neuronal processes, these methods lack the optimized performances with lower degree of accuracy to fully capture or decode complex behavior [3–5]. Machine learning (ML) offers neural decoding tools with potential in significantly boosting performance in network analysis, and thus provide deeper insights into how neuronal network activity gives rise to a behavior. Compared to traditional ML methods, advanced models such as deep learning, not only significantly improve decoding accuracy but also provide accurate identification of critical input features [6].

Brian-inspired artificial intelligence (AI), such as artificial deep neural networks, has been originated in brain networks either to decode and predict brain signals for specific behavior, or to develop innovations with AI such as brain machine interfaces (BMI). This bidirectional analysis between AI and the brain, however, has been generally limited to the data extracted from macroscale brain networks indicating brain regions (using techniques such as functional magnetic resonance imaging or local filed potentials) [7–11], from large populations of neurons (using electrophysiological monitoring such as electroencephalography) [12, 13] , or from isolated single neurons [9]. Recent advances in brain tracing and readout technologies allow real-time monitoring of detailed brain signals from individual neurons extending to the large neuronal populations. Among these, *in vivo* two-photon (2P calcium imaging offers the ability to track the activity of broad neuronal populations with high spatial resolution and in identified cell types of interest. This technology thus is a powerful tool for elucidating whether neuronal populations give rise to function and has been widely used in recent studies of the neuronal circuits underlying motor control, sensory perception and cognitive function [14–16].

Motor cortex has long been known to play a central role in the generation of complex movements, however, specific functional organizations of its subregions, at the level of neuronal populations, remain to be understood. Complex movements are composed of two fundamental features: discrete movements such as reaching that are defined as continuous movements from a start point to an end point, and rhythmic movements including locomotion, that are defined as repetitive, oscillatory movements [17, 18]. In rodent, forelimb complex movements are shown to be controlled by two distinct cortical functional regions: the caudal forelimb area (CFA) and the rostral forelimb area (RFA) [19]. Several models have suggested the specific functions for these areas or the functional relationships between RFA and CFA in complex forelimb movements [20–24]. Yet their exact functions at the level of specific neuronal populations remain controversial and not clear. Moreover, it has been demonstrated that these two areas have crucial roles in the recovery of motor function after neurological disorders such as stroke, underlining the importance in an understanding of their functions [25, 26].

In this study, we implemented machine learning models to determine whether we can predict the complex motor behavior of gait in animals on a grid-walking wheel based on neuronal network activity. Advanced 2P calcium imaging has been employed to track the activity of excitatory neuronal populations in CFA, RFA and somatosensory hindlimb (S1HL) in the head-fixed grid-walking animals. We first demonstrate that brain signals at the level of neuronal population can be used to decode complex behavior using ML models. Second, we generate a feed forward deep neural network with high decoding efficiency compared to previous models in ML. Our data demonstrate significant differences between RFA and CFA decoding efficiency in both models, indicating possible functional specificity. This result could provide insight into the principles underlying the neural code of movements and opens new routes in developing brain-inspired neural networks based on the data extracted from specific neuronal populations.

## Materials and Methods

### Experimental model

All procedures were performed in accordance with the National Institute of Health Animal Protection Guidelines. CaMKII-tTA/tetO-GCaMP6s transgenic mice were achieved from crossing B6;DBA-Tg(tetO-GCaMP6s)2Niell/J with B6.Cg-Tg(CamK2a-tTA)1Mmay/DboJ in order to have constant expression of GCaMP6s. 3 months old CaMKII-tTA/tetO- GCaMP6s males were used in this study. Mice were housed under pathogen-free conditions and were maintained on a reversed 12:12 hour light: dark cycle with free access to irradiated pellets and sterilized, acidic water.

### General surgery

Each mouse underwent a single surgery. Surgical procedures were performed under 1.5% isoflurane anesthesia and each animal was injected subcutaneously with carprophen (4 mg/kg) one hour before surgery and at 24-, 48- and 72-hours post- operative for pain relief. Craniotomy was performed in a stereotactic frame (David Kopf Instruments, CA, USA) using a hand-held dentist drill with 0.2 mm burr. The scalp was shaved, cleaned, and resected, the skull was cleaned, and the wound margins glued to the skull with tissue glue (VetBond, 3M), and a 6 mm circular craniotomy was made over the motor, premotor and somatosensory regions to make the possibility of recording from RFA, CFA and S1HL. An 8mm round coverslips (Harvard Apparatus) was then placed on the craniotomy and glued in place first with tissue glue (VetBond, 3M), and then with cyanoacrylate glue (Krazy Glue) mixed with dental acrylic powder (Ortho Jet; Lang Dental). The precis location of bregma was then labeled on the coverslips detecting and stabilizing the FOVs. A costume-made head-bar was attached to the neck of each animal for head-fixed recording. Mice were then left for one week to recover.

### Behavioral training and Forelimb tracking

mice were handled extensively before being head restrained and habituated to the grid-walking wheel. A costume-made grid walking wheel was created to track complex movement under the 2P imaging. A 1 x 1 cm squares grid was cut into 7-inch-wide sections and fit into the backs of a 9- inch-wide section. To motorize the wheel, a small DC motor was connected to the axle of the wheel such that the grid could be driven forward. This adaptation forces the mouse to move forward with adjustable velocity. Behavior was recorded using a pair of infrared-sensitive cameras that were mounted facing the mouse and at 90° to the right paw. Infrared illuminations enabled behavioral tracking while performing 2P microscopy in a darkened microscope enclosure. After the period of habituation and training, all sessions were recorded at the same velocity of the wheel. Behavioral analysis was performed manually based on previously established criteria and footsteps frames were labeled (20).

### Two-photon imaging

*In vivo* calcium imaging was performed with a Neurolabware 2P microscope (FOV: 512 x 720 µm) and pulsed Ti:sapphire laser (Chameleon, Coherent) tuned to 920 nm wavelength. The 8 kHz resonant raster scanner equated to 512 lines at 30 Hz bidirectionally and resulted in acquiring images at 15.42 Hz with a 16x objective lens (0.8 numerical aperture; Nikon). A dichroic mirror (Semrock) is used to filter the captured light and allow for imaging corresponding to GCaMP6s signals. Captured images were costume-programmed in Scanbox (https://scanbox.org/).

Custom-written MATLAB software and ImageJ (NIH) were used to analyze extracted data. Raw recorded files were converted to .tif files. Regions of interest (ROIs) related to somas with an intensity higher than 30% of the background intensity (darkest region of a video) were selected for each FOV, as described previously (19). These ROIs were also size constrained from 50 to 150 µm^2^. The output matrix was achieved with all potential ROIs and their fluorescent traces, and each ROI was manually inspected for its morphology and fluorescent traces. Fluorescence intensity was converted to dF/F0 values, which were calculated by subtracting the average fluorescence of the ROI from the current fluorescence at each frame, then dividing by the average fluorescence (F(t) – F0)/F0). Calcium transient peaks were detected by applying a MATLAB smoothing and peak detection function to the waveform. Peaks that were not greater than the root mean square of dF/F0 were not considered calcium transients. A deconvolution algorithm was then applied to all dF/F0 traces to remove the non-physiological slow decay of GcaMP6s signals and to sharpen the calcium transients (19,23).

### Neural decoding

To assess whether the neuronal activity at level of ensemble was predictive of complex movement, we used SVM and FFDNN classifiers, that we trained on instructed trials to predict footsteps. Extracted dF/F0 and deconvoluted dF/F0 were first used as sources for feature extraction. However, because of the higher decoding accuracy, only deconvolved signals were used for further analysis. Data from behavioral analysis was used as sources of target data (model prediction). For behavioral signals we used binary classification as correct right paw footsteps were considered as 1 and all other conditions including foot-faults or left paw footsteps were considered as 0. First all neurons regardless of selectivity entered the analysis, but better results were achieved with feature selection. For feature selection, the absolute value Pearson correlation coefficient (PCC) between each feature (neuron) and labeled frame (footstep) was calculated. Features with an absolute PCC higher than 0.03 were selected for SVM training. Five-fold cross validation was used to evaluate model performances, that results in evaluating the model 5 times and each time using four fifth of the data for training and one fifth for testing. Since the data was time dependent and shuffling the data can cause data leakage, we did not shuffle the data in the cross validation and kept a 10 frames distance between each neighboring folds.

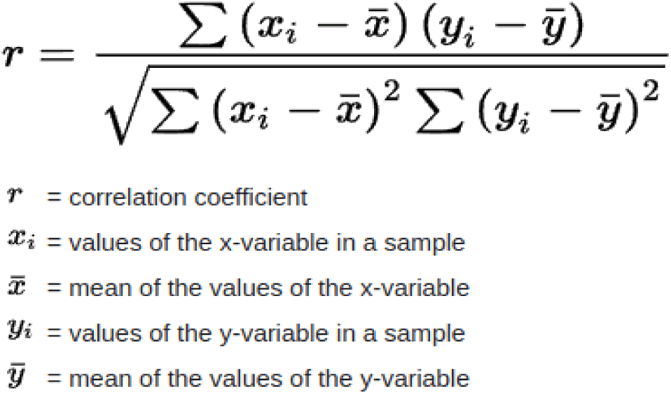

The implementation of SVM in this study was achieved from Python’s Scikit-learn library (version 1.3.0) with all parameters set to default for training classifier. The default kernel for creating boundaries was a radial basis function here.

For FFDNN, all features with deconvolved signal were selected. Deep neuronal network was modeled with Keras high-level API of the TensorFlow platform (version 2.11). Dense layer type was used for all layers of the network. Rectified linear unit (ReLU) was used for the activation function of all layers except the last layer. Apart from these main layers, we applied a dropout layer with the rate of 0.2 between the first and the second layer that led to a better regularization and prevented overfitting of the model. Activation function of the last layer was sigmoid. The Binary Cross-Entropy Loss function from Keras was used to compute the differences between predicted probabilities and actual labeled frames (footsteps).

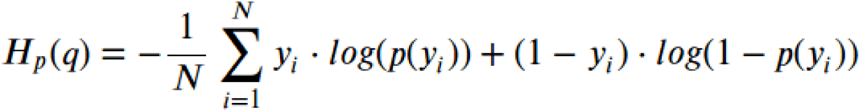

Adam algorithm was applied as the optimizer in our FFDNN. Five-fold cross validation was used to evaluate the model’s performances.

### Quantification and statistical analysis

Data analysis was performed using custom-written Python3 or MATLAB scripts (codes will be made available on request) and Prism software (GraphPad 9.5). All statistical details of experiments and analysis can be found in the main text or in the figure legends. Statistical significance was considered when p < 0.05 unless otherwise stated. Data were tested for normality and parametric/non-parametric tests were used as appropriate.

## Results

To evaluate whether a complex movement can be decoded from activity of excitatory motor cortical neurons at the level of neuronal ensembles, we developed a head-fixed grid-walking paradigm to simultaneously monitor behavioral performance and neuronal activity (Fig.1B). The grid-walking task measures forelimb placement on a challenging gird during exploratory gait and has been widely used to track complex movements and motor-skill learning, as well as deficits in descending motor pathways in brain diseases such as stroke or traumatic brain injury [27]. Animal was trained on the wheel and complete footsteps, representing complex movements, were evaluated. Each footstep is composed of continuous movement from the start point (indicating releasing the paw from the initial grid) to the end point (indicating grasping the next grid) (Fig.2B). Since the animal was head-fixed, the setup was motorized to generate rhythmic movements representing repetitive footsteps with identified velocity. Trained animals exhibit an increase in the number of complete footsteps and decreased number of foot-faults that reached a stable level after 4 days of training (Fig.1C). To subsequently track neuronal activity in different brain regions, CaMKII-tTA/tetO-GCaMP6s transgenic mice were employed. This line expresses GCaMP6s broadly in excitatory neurons [28]. An 8 mm cranial window was stereotaxically located on the head of animal to provide the possibility of recording from layers II/III of contralateral CFA, RFA and S1HL in individual animals. These layers represent a high number of intra-cortical neurons, which would be involved in information processing vs corticospinal or other cortical output functions [29–32]. Recording from S1HL was used as a control in this study [33]. 3000 frames, at 15.32 Hz, were recorded from each region using 2P imaging and in parallel, two CCT cameras were used to track movement of head-fixed animal (Fig.1 & 2). The complete footsteps from right paw were evaluated and 2P imaging frames corresponding to each footstep were labeled. For each footstep, the initial frame was labeled as t_0_ and the end frame was indicated as t_0_+x. The majority of footsteps in our behavioral analysis paradigm are composed of 5 frames (t_0_+4) (Figs. 2B), however, footsteps with different duration were also considered. Recording sessions were collected across 8 days at a 2-day interval and calcium transients for each region of interest (ROIs, representing neurons) were extracted using previously established image processing methods (ref.). Time traces of calcium signals from each ROI were converted to dF/F_0_ values as (F(t) – F_0_)/F_0_ (Fig. 2A). Calcium signals have a long-decay constant that is a property of GCaMP rather than the neuronal signal. Thus, for further analysis, we deconvolved calcium transients to infer the time course of instantaneous spiking rate changes [26, 34]. During the labeled frames (footsteps), Layers II/III excitatory activity was heterogeneous across the sampled neuronal subsets and temporally sparse, however, some neurons displayed calcium transients that linked to paw movements during footstep (Fig. 2B). We first computed the mean dF/F_0_ of each neuron in a given neuronal network. Averaged dF/F_0_ across all neurons for each network related to each frame did not show significant differences for labeled versus non-labeled frames. We thus evaluated the activity of individual neurons related to labeled frames; each footstep was considered as R (0 for no footstep and R=1 for a complete footstep) and the correlation coefficients (CC) between individual neurons and R were calculated. Neurons with CC values higher than 0.03 were used for ML algorithm training. It should be noted that other cutoff values for correlation coefficient have been applied as well, however, the best output achieved with this 0.03 cutoff. Neurons with CC higher than cutoff (threshold) were classified as movement responsive versus non-responsive neurons with CC lower than threshold (Fig. 4A-B). Interestingly, the ratio of responsive vs non-responsive neurons was distinct for each brain region and a higher ratio of responsive neurons was detected in CFA compared to RFA recorded networks (357 of 518 neurons, mean = 74.52% for CFA and 218 to 421 neurons, mean = 36.82% for RFA) (Fig. 4A-B). Each included neuron (responsive) was considered as a feature (input) resulting in a high dimensional data (input). Because of the high dimensionality of input, a support vector machine (SVM) model was first used for prediction of movement based on calcium signals. Non-linear classification was trained from both deconvoluted and non-deconvoluted calcium signals. Since deconvoluted input demonstrated significantly better prediction of footstep for all brain regions (data not shown), only deconvoluted signals were used for further analysis. For each recording session, representing an excitatory neuronal network, we then evaluated F-score as a measure of our model’s accuracy as [(precision x recall) / (precision + recall) (Fig. 3B). Given the achieved high F-score decoding, our results demonstrated that layers II/III excitatory neuronal responses (signals) at the level of ensembles could provide robust movement-related information that can be used for movement decoding (Fig. 3B). In addition, distinct decoding accuracy was observed; CFA exhibited a significantly higher F-score compared to RFA (CFA SVM mean = 75.1% and RFA SVM mean = 66.2%, p = 0.0004) (Figs. 4A-B). Considering the higher ratio of responsive neurons in CFA, as well as higher CFA decoding accuracy, the results suggest a dependency on neurons with high decoding accuracy and based on brain region. We then built a feedforward deep neural network (FFDNN) of 5 layers that connects the inputs, indicating deconvoluted calcium signals, to the sequential layers of hidden units, and then connects to the output, indicating movement decoding. Our input layer is composed of 512 neurons, connecting to hidden layers with 64, 8 and 4 neurons respectively, and lastly the output layer with a single neuron (Fig. 3A). The layers of our FFDNN have been sequentially connected to each other via linear mappings followed by nonlinearities [35]. Evaluating F-score, our data demonstrated an even higher level of decoding accuracy with FFDNN compared to SVM in both CFA and RFA (CFA FFDNN mean = 81.243, p = 0.0094 and RFA FFDNN mean = 71.733, p = 0.0195). Interestingly, like SVM, we observed a significant difference between CFA and RFA F-scores achieved from FFDNN (CFA mean = 81.243% and RFA mean = 71.733%, p = 0.0002). This data could suggest that movement-specific (here, footstep) information flow is routed through a selective ensemble of layers II/III excitatory neurons with higher distribution among CFA neuronal networks (neurons).

**Fig.1.**
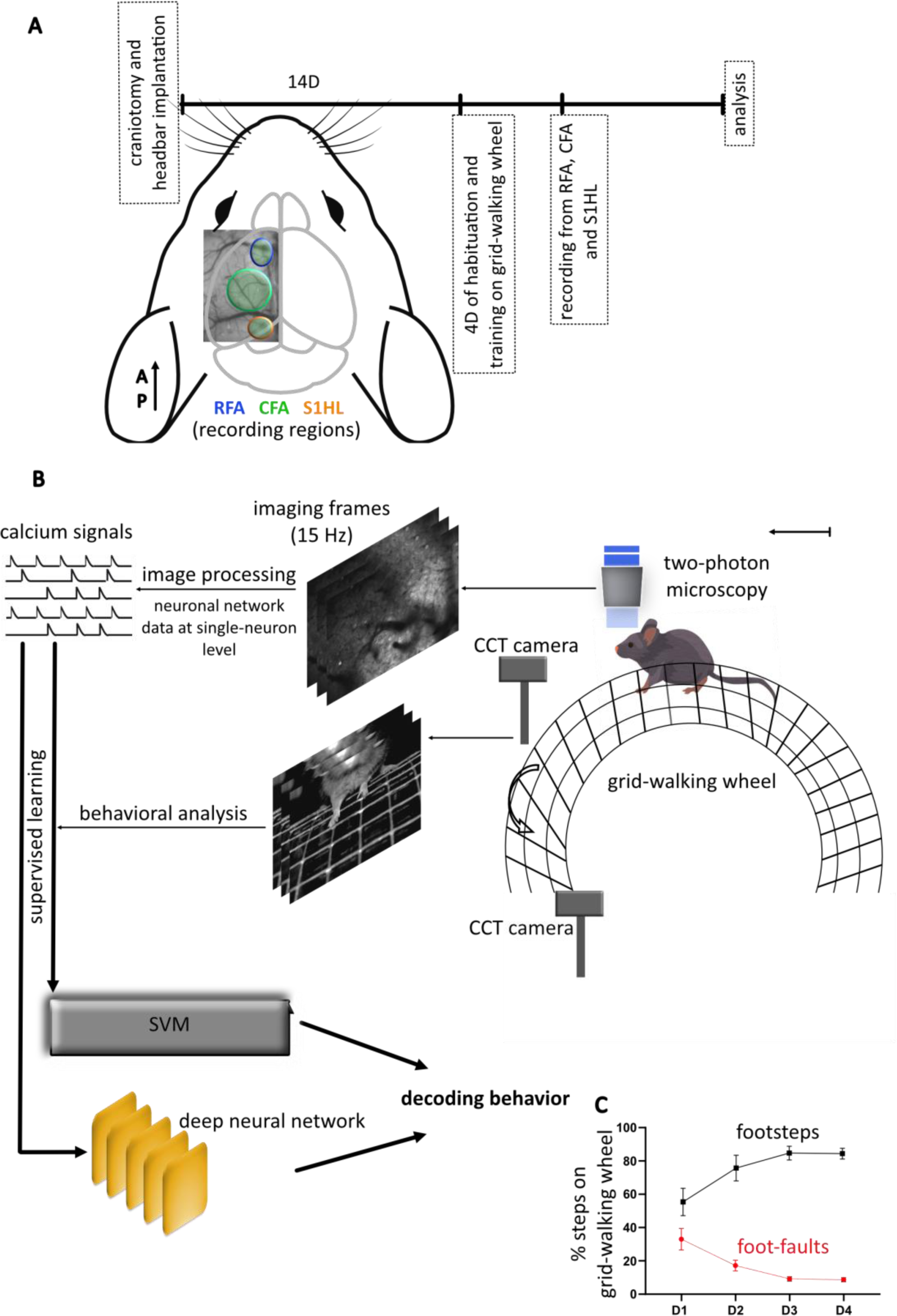
Experimental design. **A)** Timeline of experiment: 2 weeks after cranial window surgery (skull and dura removing), teto-GCaMP6s males are habituated and trained on the grid-walking wheel for 4 days and with 15 minutes duration. Recordings from CFA, RFA and S1HL are performed for each head-fixed moving mouse. Image of left hemisphere cranial window over regions of interests in green circles (each region has color code, indicated in 1B). **B)** Representative image of experimental setups including two-photon microscopy and grid-walking wheel. A series of image processing algorithms are used to extract the calcium signals from two-photon recordings. Two CCT cameras are used to record behavioral performance. Behavioral data are analyzed, the frames are labeled for footsteps. Supervised machine learning approaches, a feed forward deep neural network and a support vector machine (SVM) are applied to decode behavior from calcium signals. **C)** Habituation on the grid-walking wheel for 4 days indicates increased number of footsteps and a decreased number of foot-faults that reach to the stable level after 4 days. CFA indicates caudal forelimb; RFA for rostral forelimb; S1HL for somatosensory hindlimb. Scale bar: 25 µm.

**Fig.2.**
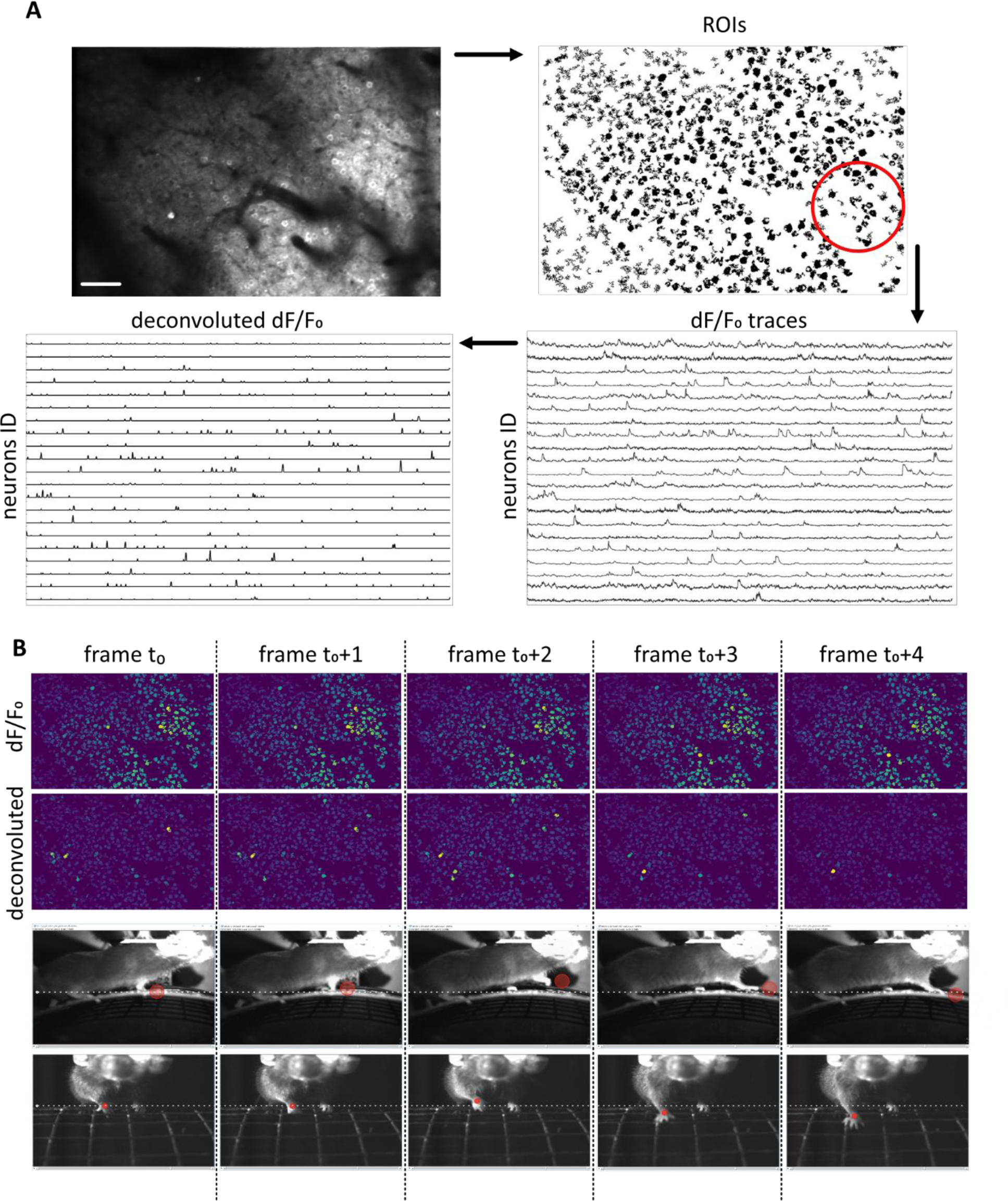
Neuronal network activity and behavioral analysis. **A)** Upper left: average of 3000 frames displaying CFA excitatory neurons from one field of view. Upper right: ROIs detected after segmentation processing on the video recorded from left image. Lower right: example of extracted dF/F_0_ traces for indicated neurons in red circle, each neuron has specific ID. Lower left: example of traces after applying deconvolution algorithms on the right image dF/F_0_. **B)** The mouse is required to learn and perform a complex behavior of complete footsteps that includes reaching and grasping the grid wire, from the start point (t_0_) to the end point (t_0_+x, here x=4) (discrete movement) and adapt to perform repetitive footsteps on the running motorized wheel with identified velocity (rhythmic movement). The upper panel represents distribution of ROIs that are achieved based on dF/F_0_ or deconvoluted dF/F_0_ measurement. Each upper frame corresponds to lower behavioral frame. Lower images include CCT recordings from two angels for accurate behavioral analysis. Red circles indicate the position of right paw that is used to determine footstep. Scale bar: 50 µm.

**Fig.3.**
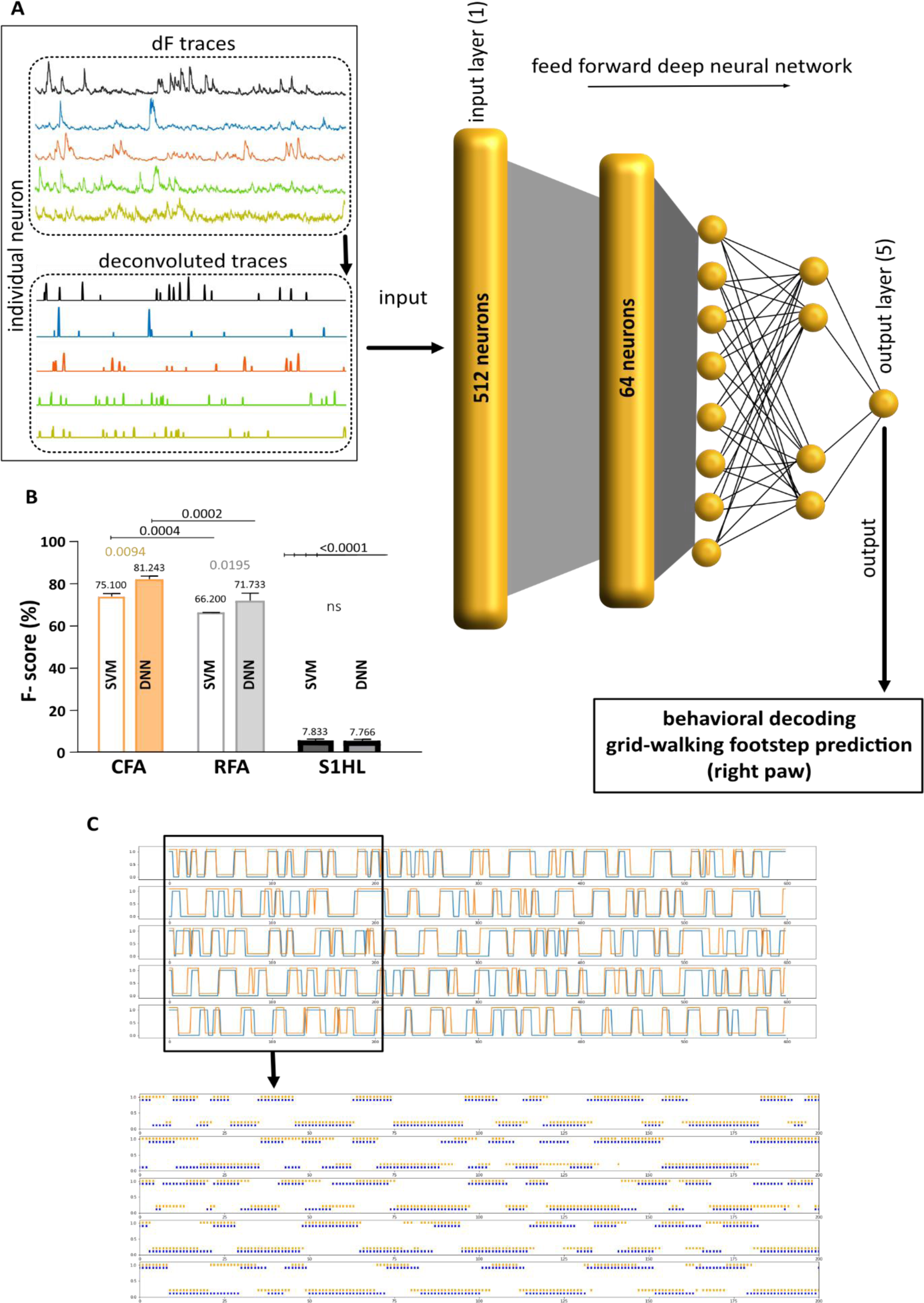
Decoding complex movement in CFA and RFA networks. **A)** Schematic of deep neural network architecture: deconvoluted signals extracted from each ROI in two-photon microscopy are used as neural modalities representing input. Deconvoluted signals are calculated for each single neuron. The feed forward deep neural network (FFDNN), developed here, includes five layers with input, hidden layers, and output layer; input layer-1 composites of 512 neurons and accordingly 64, 8 and 4 neurons for the following layers. The output layer-5 has a single neuron. The output of the FFDNN is prediction of footstep. **B)** Decoding accuracy is quantified by measuring F-score between true and decoded footsteps across all trials including 3 recoded movies per animal and for each brain region applying both DNN and SVM models. C) Example of DNN decoding within a given acquisition. The upper panel shows the network prediction of footsteps in blue and the true footsteps in orange achieved from 600 frames in CFA. The lower panel corresponds to extracted 200 frames from the upper image, indicating true (orange) and predicted (blue) footsteps in more detail. Each dash represents a single frame. One-way ANOVA multiple comparisons, Tukey’s multiple comparisons test were used for statistical analysis (2 mice and 8 sessions total per brain region). Statistical significance was considered when p < 0.05 and was indicated in B.

**Fig.4.**
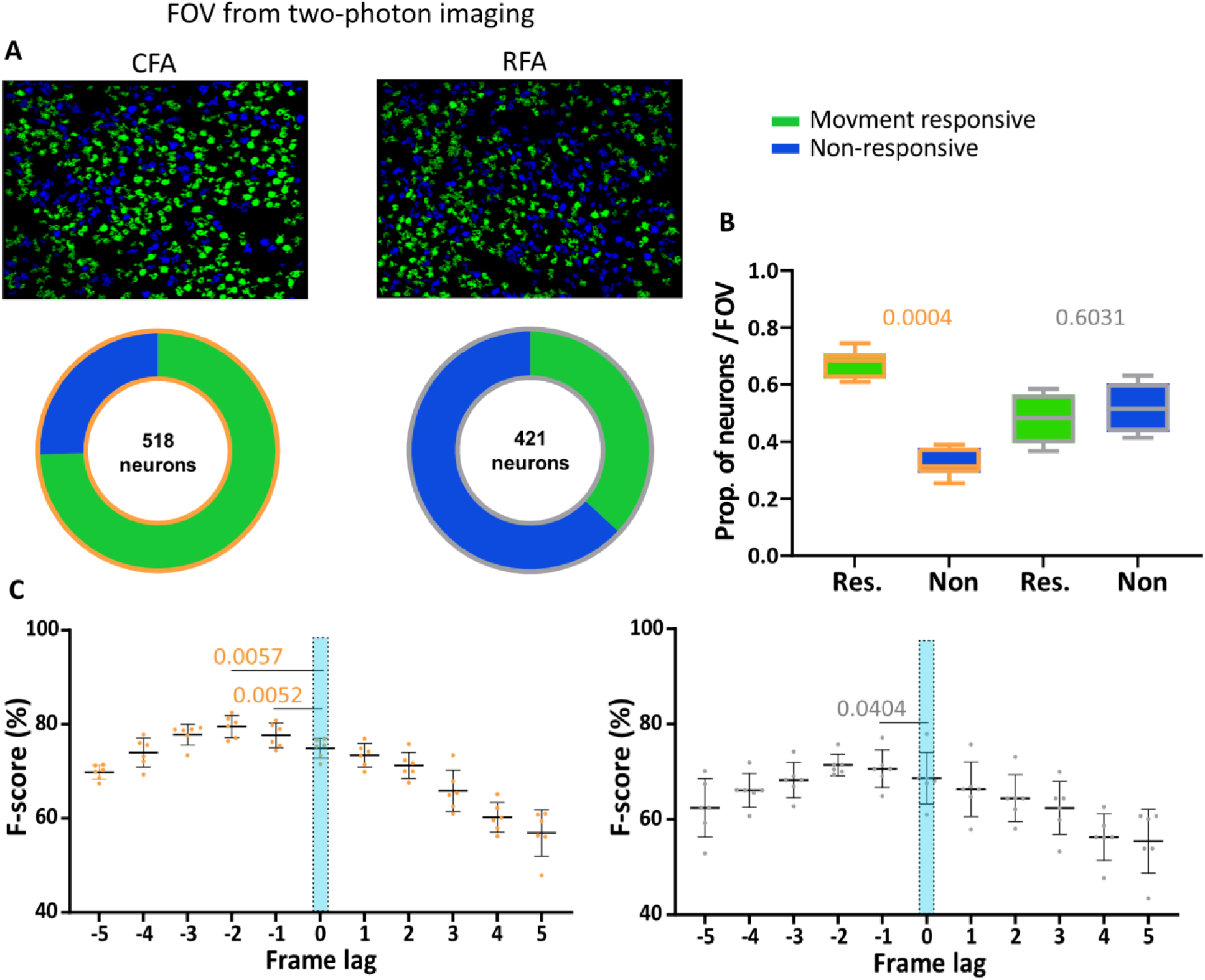
Functional dissociation of spatiotemporally concurrent neuronal ensembles in CFA and RFA networks. **A)** top: example of a field of view from 2p recording in CFA and RFA networks. Green shows movement responsive neurons and blue indicates non- responsive neurons in FOV for SVM models. The lower panel demonstrated the ratio of responsive and non-responsive neurons for entire experiments in CFA and RFA networks. **B)** Distinct proportion of neurons (responsive vs non-responsive) per FOV was indicated for CFA and RFA networks during movement decoding. A paired t-test was used to compare movement responsive vs non-responsive neurons. **C)** F-score computing at different frame lags before and after onset of movement. Each frame indicates 65 ms duration. Ordinary one-way ANOVA was used to compare F-score at various frame lags (2 mice and 8 sessions for each brain region). Statistical significance was considered when p < 0.05 as was indicated in B and C.

Several studies suggested that both CFA and RFA are involved in movement planning and execution [36]. In parallel, it has been shown that neuronal signals in the motor cortex exhibit delays, in conduction and processing, to reach the muscles of interest and execute movement. The decoding accuracy of movement trajectories, thus, can be amplified by neuronal subpopulation signals at the time points prior to the onset of a movement. In fact, previous studies demonstrated temporal-delay decoding for movements in which the preceding time of neuronal signals suggested for neural decoding were ranging from 100 ms to 1 s [6, 33]. We trained our deep learning models in a time decoding scheme, from 0 ms, corresponding to concurrent decoding, to 325 ms (5 frames) before and 325 ms (5 frames) after movement onset (t_0_, Fig. 2B) (Fig. 4C). The interval used here was 65 ms, indicating the temporal resolution of 2P recording. CFA neuronal networks demonstrated highest level of decoding accuracy with 65 and 130 ms long preceding time compared to the concurrent decoders (frame lag -1 (65 ms prior to movement), p = 0.0052, and frame lag -2 (130 ms prior to movement, p = 0.0057) (Fig. 4C). In RFA excitatory neuronal networks, the best decoding accuracy was only achieved using 65 ms preceding time compared to the concurrent decoding (frame lag -1, p = 0.0404) (Fig. 4C). These data suggest that the preceding time intervals could carry important movement-related information and have distinct temporal differences in CFA and RFA neuronal networks.

## Discussion

Decoding behavior from brain signals is one of the main goals in neuroscience. In recent years, machine learning models have been employed to fulfil this goal. In parallel, the brain signals have been used to generate successful transfer of biological insights into development of new brain-inspired AI algorithms. This reciprocal interaction, however, relies on the invention of novel brain tracking technologies. For the past decades, rapid progress in neuroimaging techniques have provided enormous amount of data from different scales of brain networks ranging from micro- to macroscales. Here, we used advanced *in vivo* 2P calcium imaging on the head-fixed grid-walking mice to target CFA and RFA neuronal subpopulation networks that provided the possibility of monitoring single neuron activity and then connecting this activity into the large neuronal assemblies during complex movement performance. Employing two different decoding strategies- a SVM classifier and a FFDNN, we demonstrated that neural signals from excitatory cortical neuronal ensembles could provide essential information for forelimb movement decoding. Our data suggested distinct patterns of activity-induced behavior in CFA and RFA networks. The data achieved from this work can make contributions to the field of motor neuroscience as well as brain-inspired artificial intelligence.

In rodent motor cortex, CFA and RFA are major actors in orchestrating the control of complex forelimb movements [37, 38]. We have developed a behavioral paradigm of motorized grid-walking wheel test to evaluate footsteps consisting of both locomotion like and reaching-grasping movements. This test has been widely used to evaluate motor behaviors of the forelimb during gait and exploratory rearing in several brain diseases including stroke [27]. To our knowledge, this is among first studies that characterize the performance of both CFA and RFA subpopulation networks for an identical complex movement. Our decoding strategies were focused on the accuracy of decoding. First, we found that a DNN model has significant higher level of decoding accuracy for both RFA and CFA networks than SVM classifier. Our analysis demonstrated that DNN discovered a larger number of important input features (here, neurons) than a traditional SVM decoder, suggesting that the higher decoding accuracy achieved by extracting features that are related to forelimb movements in complex and nonlinear manners. Furthermore, this work supports the notion that neural activity at fine microscale and/or ensemble level could predict behavior with high decoding accuracy through deep learning.

Recent studies demonstrating similarities in the effects of CFA and RFA stimulations on motor performances establish that these two areas are part of a highly integrated computing unit [21, 33]. We confirmed that both RFA and CFA are involved in planning and execution of skilled forelimb movement. However, our data suggested distinct patterns of activity-induced movement for CFA and RFA networks. It has been suggested that a population of neurons in a motor cortical area encode specific complex movements [39]. Instead of using different loss-of-function strategies to compare the relative contribution of CFA and RFA to forelimb complex movement, we implemented innovative ML models that allowed us the generation of predictions about excitatory neuronal networks that are otherwise difficult to compute due to their invariant responses to movement. Based on the signal extracted from 2P recording, ML models enabled the functional dissociation of spatiotemporally concurrent neuronal ensembles. Elucidating the computations in these cortical areas, we compared the neuronal activity ratio in CFA and RFA networks for purposeful complex movement. We demonstrated that CFA contained a higher number of movement responsive neurons than the RFA, indicating that majority of active excitatory neurons in CFA networks (FOV) were involved in motion planning and execution. It has been proposed that CFA and RFA receive reciprocal functional projections, generating corticocortical pathways involved in the transmission of forelimb inputs that are processed fist by CFA [40–42]. Examining the timing of activity of individual neurons with ML models, we observed that as a population, the activity of movement-related excitatory neurons diverged from baseline at a timing earlier for CFA (compare to RFA) before the movement onset and continued throughout the duration of movement.

This work provides the possibility of computing individual neuronal activity within brain areas where subpopulations dedicated to specific movements are intermingled. Our developed ML models enabled the deciphering of neural code for a complex movement that is widely used in motor neuroscience research. The knowledge achieved from this work about the role of specific neuronal ensembles can be incorporated into the design of modern brain machine interface to gain better control of external device as well as it can make contributions to the field of motor neuroscience.

## Author contribution

S.L., and S.T.C. designed research; S.L. performed research and analyzed 2P data; S.L.,

M.P. and R.L. developed neural networks; S.L., and J.C. analyzed behavioral data; S.L., and S.T.C. wrote the paper.

